# Starve, stress, and mate: Activation of the CCAAT-Binding complex overrides sterility of mating-type heterozygous *Candida albicans*

**DOI:** 10.64898/2026.11.05.764546

**Authors:** Lasse van Wijlick, Kontxi Martinez de San Vicente, Natacha Sertour, Laxmi Shanker Rai, Salomé LeibundGut-Landmann, Sophie Bachellier-Bassi, Christophe d’Enfert

**Affiliations:** Institut Pasteur, Université Paris Cité, INRAe USC2019, Inserm U1359, Unité Biologie et Pathogénicité Fongiques, Paris, France; Institut of Microbiology, Heinrich Heine University Düsseldorf, Düsseldorf, Germany; Institute of Medical Microbiology and Hospital Hygiene, Heinrich Heine University, Düsseldorf, Germany; University of Zürich, Vetsuisse Faculty, Section of Immunology, Zürich, Switzerland; University of Zürich, Institute of Experimental Immunology, Zürich, Switzerland; Medical Research Council Centre for Medical Mycology at the University of Exeter, Department of Biosciences, Faculty of Health and Life Sciences, Exeter, United Kingdom

## Abstract

Sexual reproduction in the human fungal pathogen *Candida albicans* is typically restricted by a strict genetic lock: the **a**1-α2 repressor present in standard mating-type heterozygous (*MTL**a**/α*) diploid strains, which actively represses mating competency. While severe environmental pressures are known to influence fungal development, the molecular mechanisms that allow cells to override this genetic sterility block remain poorly understood. Here, we identify the highly conserved CCAAT-binding complex (CBC) as the critical developmental hub that integrates environmental stress to unlock mating in heterozygous diploids. By combining proteomic mapping (ChIP-SICAP) and transcriptomics (RNA-Seq) to define the regulatory network of the core CBC subunit Hap5, we demonstrate that CBC hyperactivation completely overrides the **a**1-α2 repressor. This activation triggers a robust white-to-opaque phenotypic switch and initiates a mating-competence program in MTL**a**/α strains genetically locked in a sterile state. Crucially, this developmental transition is antagonized by the iron-regulatory partner Hap43, but is strictly dependent on physical interaction at the chromatin level with Cap1, the master regulator of the oxidative stress response. Our findings reveal that the CBC and Cap1 couple metabolic starvation (glucose and iron limitation) and oxidative stress to a core reproductive program, providing a mechanistic basis for how *C. albicans* bypasses genetic constraints on sexual reproduction to generate genetic diversity under hostile host conditions.

**Author Summary:** *Candida albicans* is a prevalent human fungal pathogen that constantly adapts to fluctuating environments. It can undergo a dramatic shape-shifting process called the “white-to-opaque” switch, which is strictly required for sexual reproduction. However, most natural *C. albicans* strains possess a genetic lock that keeps them in a sterile, white state. Here, we investigated the CCAAT-binding complex, a conserved group of proteins known to manage cellular metabolism and iron levels. By mapping its gene targets and protein network, we discovered that this complex also acts as a developmental switch. We found that overactivity of this complex overrides the genetic lock, allowing normally sterile cells to switch to the mating-competent opaque state. Crucially, this transition requires a physical interaction with Cap1, a key regulator of the oxidative stress response. Our findings reveal that hyperactivation of the metabolic and stress-sensing machinery can trigger sexual development in *C. albicans*, providing a potential molecular mechanism by which the fungus might unlock reproduction to generate genetic diversity when encountering environmental stress.

## INTRODUCTION

*Candida albicans*, a major opportunistic human pathogen, possesses a remarkable ability to adapt to diverse host environments, a trait largely dependent on its capacity for morphological and phenotypic plasticity. A striking example of this plasticity is the white-opaque switch, a reversible, epigenetic transition between two distinct cell types. White cells are round and form dome-shaped colonies, while opaque cells are elongated, metabolically distinct, and form flatter colonies. Crucially, this switch is directly linked to sexual reproduction, as opaque cells represent the mating-competent form of the fungus [1,2,3].

Mating in *C. albicans* is tightly controlled by the mating-type locus (*MTL*). The vast majority of clinical isolates are diploid and heterozygous at this locus (*MTL***a**/α), a genetic state that actively represses mating [4,5]. In these cells, the proteins **a**1 and α2, encoded by the respective *MTL* alleles, form a heterodimeric repressor (**a**1-α2) that blocks the expression of genes required for both the opaque state and mating [6]. Consequently, mating and the prerequisite switch to the opaque form are typically restricted to cells that have undergone loss-of-heterozygosity and become homozygous or hemizygous at the *MTL* locus (*MTL**a**/**a***, *MTL**a**/Δ*, *MTLα/α* or *MTLα/Δ*). While mating canonically occurs between cells of opposite mating types, *MTL*-homozygous opaque cells can also undergo same-sex mating (e.g., **a**/**a** × **a**/**a**). Recent studies have highlighted that severe environmental cues, particularly glucose starvation, can influence the white-to-opaque transition and promote same-sex mating, suggesting a profound link between the metabolic status of the cell and its developmental potential [3,7,8].

Cellular adaptation to such environmental signals is governed by intricate transcriptional networks. A central player in these networks across eukaryotes is the CCAAT-binding complex (CBC), a highly conserved transcription factor that recognizes the CCAAT motif in the promoters of numerous genes [9]. The core of this complex is a heterotrimer composed of the Hap2, Hap3, and Hap5 subunits. In *Saccharomyces cerevisiae*, the CBC is a critical activator of respiration, and null mutations in its core subunits render cells unable to grow on non-fermentable carbon sources [10]. In contrast, the CBC in pathogenic yeasts often exerts dual functions. For instance, in *Candida glabrata*, the CBC acts as both an activator of respiration and a repressor or activator of iron homeostasis genes depending on its competitive association with the accessory subunits Hap4 or Yap5 [11,12]. Similarly, the *C. albicans* CBC acts as both an activator and a repressor [13]. The *C. albicans* genome encodes two Hap3 paralogs, Hap31(orf19.517) and Hap32 (orf19.4647), which exhibit distinct functions and variable redundancy depending on the specific stress condition [14,15]. Beyond core metabolism, the *C. albicans* CBC is a master regulator of iron homeostasis, functioning in concert with the accessory subunit Hap43 to regulate iron uptake and utilization genes during iron deficiency [16]. Reflecting its pleiotropic nature, the functional repertoire of the *C. albicans* CBC has recently been expanded to include the regulation of cell wall integrity and biofilm formation via the Rhb1-Mkc1 signaling pathway [15]. Consistent with these diverse roles, mutants lacking CBC subunits in *C. albicans* exhibit profound defects in morphogenesis, oxidative stress adaptation, and cell wall remodeling.

While the CBC is known to be a crucial regulator of metabolic and stress-related processes, the full scope of its regulatory network -and the mechanisms by which it integrates diverse host-derived signals -remains incompletely understood. To decipher how this complex orchestrates environmental adaptation, we comprehensively mapped the regulatory landscape of the core subunit Hap5 using a combination of transcriptomics (RNA-seq), genome-wide binding profiling (ChIP-seq), and targeted isolation of chromatin-associated proteins (ChIP-SICAP, [17,18]). This multi-omics approach confirmed the CBC’s canonical role in orchestrating carbohydrate and iron metabolism, but unexpectedly revealed a striking transcriptional signature normally associated with mating-competent cells, even in *MTL**a**/α* heterozygotes. We demonstrate that overexpression of *HAP5* overcomes the canonical **a**1-α2 repressor to induce robust white-to-opaque switching and enable successful mating. We further show that this novel developmental function is antagonized by the iron-regulatory partner Hap43, but absolutely requires physical interaction with Cap1, the master regulator of the oxidative stress response. This work establishes the CCAAT-binding complex as a molecular hub that couples metabolic and oxidative stress sensing to a core developmental program, providing a mechanism for *C. albicans* to bypass genetic barriers and engage in sexual reproduction under hostile environmental conditions.

## RESULTS

### Transcriptomic and chromatin-binding profiling defines the core Hap5 regulon

To comprehensively map the regulatory network of the CCAAT-binding complex (CBC) in *C. albicans*, we combined genome-wide chromatin immunoprecipitation (ChIP-seq) with transcriptomic analysis (RNA-seq) of *C. albicans* strains engineered to overexpress wild-type or TAP-tagged Hap5 from the P*_TET_* promoter, as this approach has proven successful for several other transcription factors [19,20]. Binding sites of the core subunit Hap5 were identified using a modified ChIP-SICAP approach with a TAP-tagged Hap5 construct, which allows rigorous washing to drastically reduce protein background [17,18] **(Fig. S1)**. Using the bPeaks program [21], a total of 1039 significant Hap5 binding sites were identified, which were mainly located in the upstream regulatory regions of ORFs across all 8 chromosomes. The transcriptome was analyzed by RNA-seq after 2 hours and 4 hours of tetracycline-induced *HAP5* overexpression in rich medium. Transcriptional changes included 213 upregulated and 382 downregulated genes at the 2 h time point, and 300 upregulated and 383 downregulated genes at the 4 h time point (fold-change ≥1.5 or ≤ -1.5, Padj < 0.05; **Table S2**). The overall concordance with genome-wide binding was strikingly high: more than 85% of all obtained binding sites could be assigned to at least one gene with altered expression **(Fig. 1A)**, indicating that Hap5 acts directly as both an activator and a repressor of gene expression, consistent with observations for its homologs in *S. cerevisiae* [22] and *C. glabrata* [11].

**Figure 1.**
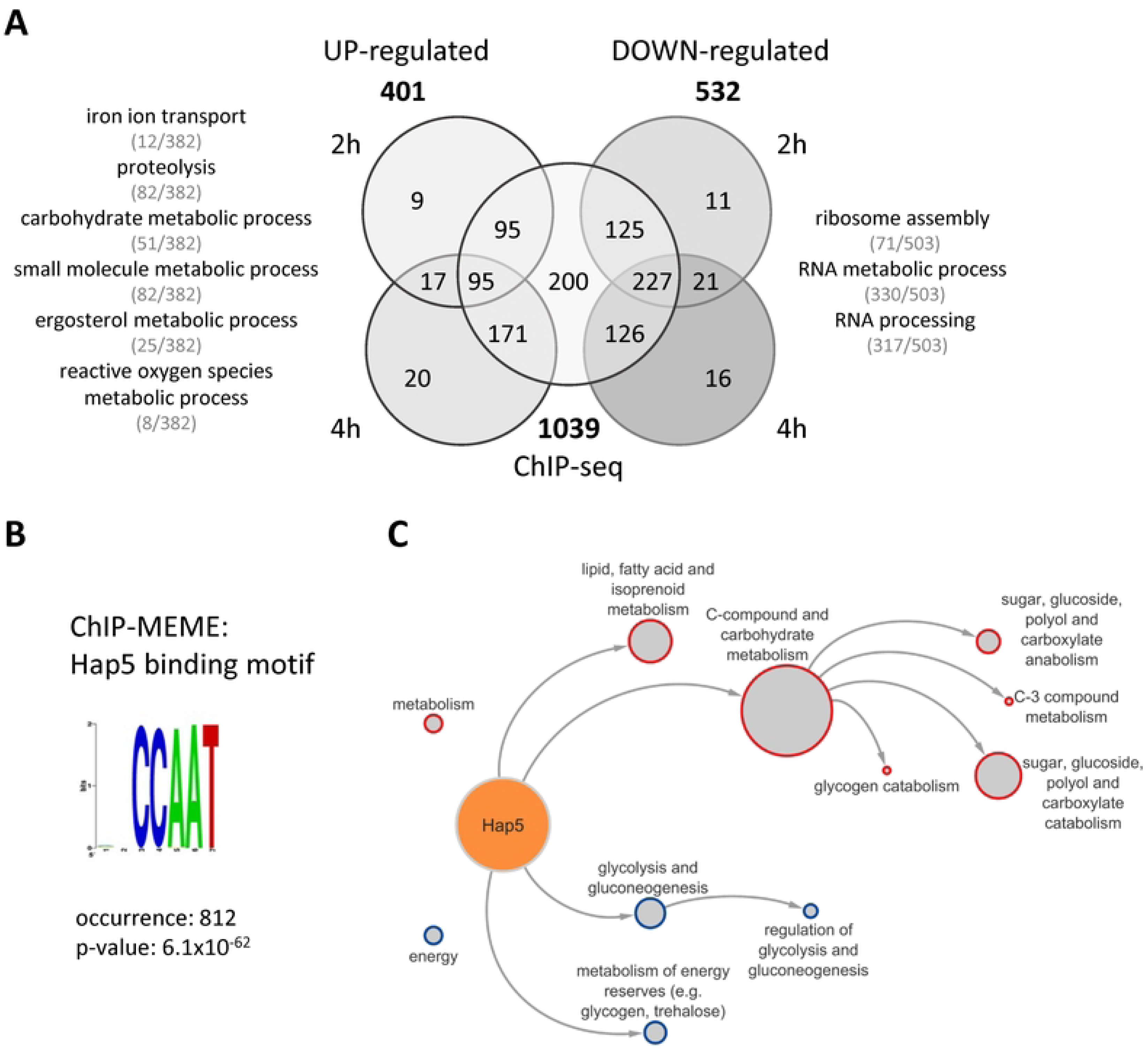
Transcriptomic and chromatin-binding profiling defines the core Hap5 regulon. Combination of RNA-seq and ChIP-seq was used to establish direct and indirect target genes of Hap5 in YPD medium at 30 °C. **(A)** Venn diagram showing the overlap of genes differentially expressed in response to *HAP5* overexpression (after 2 h and 4 h of induction) and genes possessing a Hap5-TAP binding event in their upstream regulatory sequence. Gene Ontology (GO) analysis was used to identify functional categories of the enriched gene sets (selected categories are shown). **(B)** Identification of the highly conserved Hap5 binding motif using ChIP-MEME. **(C)** Functional categories related to metabolism and energy are presented as a network. The size of the nodes is proportional to the number of associated genes belonging to the respective term. Genes in both categories were highly upregulated in strains overexpressing *HAP5*.

Gene ontology (GO) analysis revealed that upregulated Hap5 target genes are highly enriched for processes involved in carbohydrate metabolism, ergosterol metabolism, iron ion transport, and proteolysis. Conversely, downregulated targets are predominantly involved in ribosome assembly and RNA metabolic processes **(Fig. 1A, Table S3)**. Motif enrichment analysis of the Hap5 binding sites using ChIP-MEME [23] identified a highly conserved CCAAT-motif **(Fig. 1B)**, as expected based on homology to *S. cerevisiae* [24,25]. A closer examination of the metabolic targets revealed that Hap5 positively regulates glycolysis, energy production from glycogen and trehalose, as well as lipid and fatty acid metabolism, as do other CBC components **(Fig. 1C)**, underscoring the central role of this complex in fungal energy management.

### Regulation of iron-responsive genes and virulence effectors by Hap5

The best-characterized co-regulator of the CBC in *C. albicans* is the Hap43 protein that modulates the cellular response to iron deficiency [16,26]. To understand how Hap5 functionally integrates with iron homeostasis, we specifically analyzed the expression of genes encoding components of iron acquisition **(Fig. 2A)**. Under iron deficiency [27], the expression of iron-responsive regulators shows an induction of the CBC components (*HAP2*, *HAP31*) and *HAP43*, while the antagonist *SFU1* is strongly repressed. *HAP5* overexpression mirrored this physiological response to a large extent: expression of the negative regulator *SFU1* was initially inhibited, consistent with feedback regulation [28], and components of heme acquisition (*RBT5*, *PGA7*, *CSA2*) were significantly upregulated.Strikingly, genes directly bound by Hap5 (except *CFL4* and *CFL5*) showed upregulation upon *HAP5* overexpression similar to that observed in iron-deprived cells, suggesting targeted overexpression of *HAP5* mimics an iron deficiency-like transcriptional response.

**Figure 2.**
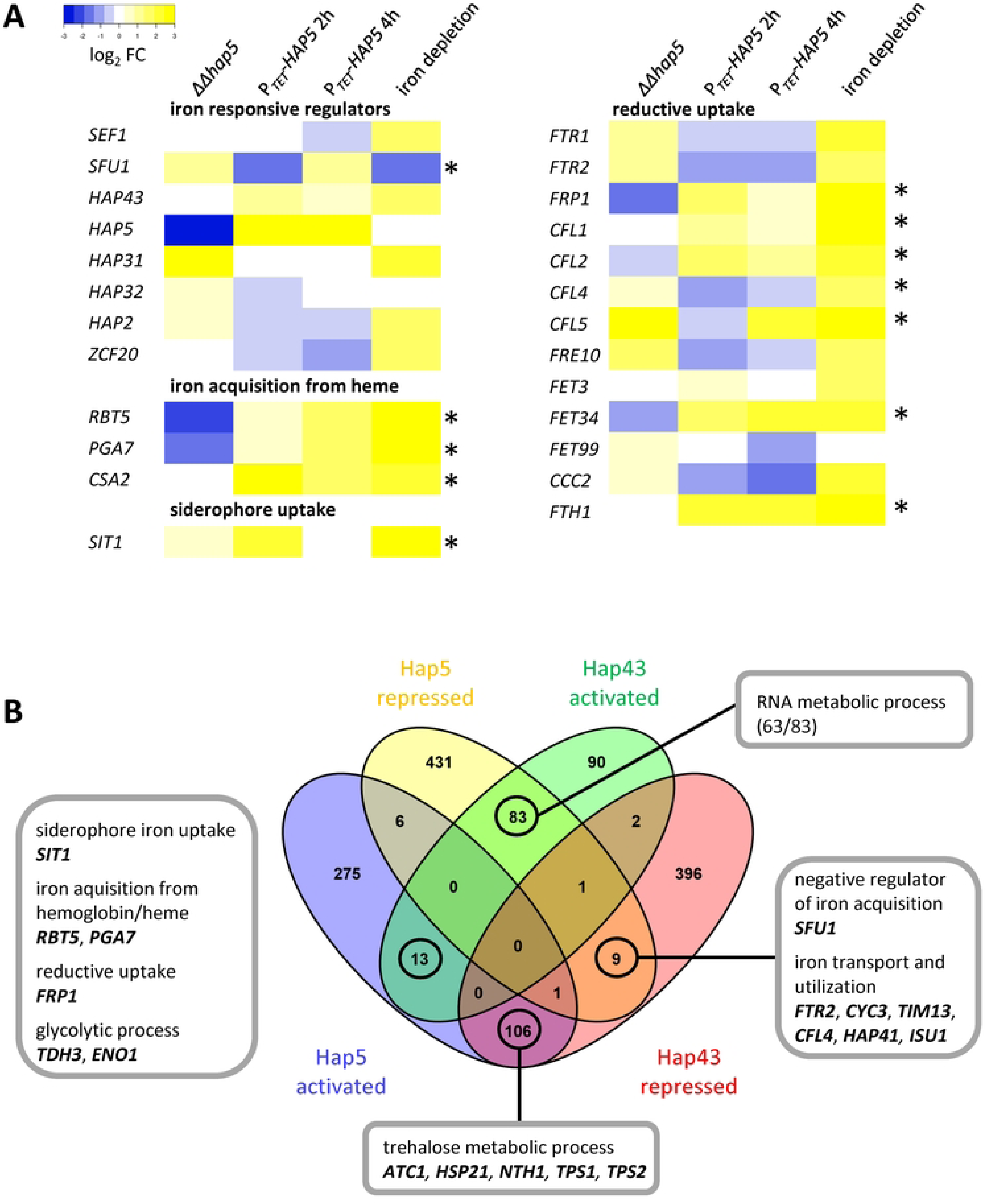
Regulation of iron-responsive genes by Hap5 and Hap43. **(A)** Heat map showing log2 fold-change levels of selected transcriptional regulators involved in iron regulation and genes involved in iron acquisition. Transcriptomic data of wild-type cells incubated under iron deficiency were retrieved from [27] for comparison. The asterisk (*) indicates direct binding of Hap5 to the upstream regulatory sequence of the indicated gene. **(B)** Venn diagram showing the overlap of identified Hap5 target genes and Hap43 target genes. Hap43 target genes were retrieved from [16].

To determine how Hap5 interacts globally with the known iron regulator Hap43, we subsequently compared their broader regulatory networks **(Fig. 2B, Fig. S2, Fig. S3)**. Surprisingly, a direct comparison showed complex, often antagonistic dynamics. For instance, genes involved in RNA metabolism or trehalose synthesis that are repressed by Hap43 appear to be activated by Hap5, and vice versa, indicating that while both transcription factors cooperate in the regulation of specific iron acquisition processes, Hap5 also exerts distinct, Hap43-independent regulatory functions.

Beyond canonical iron transporters, transcriptome analysis revealed that Hap5 regulates genes encoding secreted aspartyl proteinases (*SAP1-10*, *SAP98*, *SAP99*) and the candidalysin toxin (*ECE1*), which are important for host tissue invasion and could represent an alternative mechanism for iron acquisition via host protein degradation [29,30] **(Fig. S4A)**. However, phenotypic assays assessing host cell damage, proteolytic, and phospholipase activity showed that both *HAP5* overexpression and *hap5* deletion resulted in significantly reduced host damage and enzymatic activity *in vitro* **(Fig. S4B-D)**. This suggests a tightly balanced, context-dependent role for Hap5 in mediating the expression of virulence effectors.

### The transcript profile of strains overexpressing *HAP5* displays a *MTLa* signature

While exploring the *HAP5*-overexpression transcriptome for broader developmental impacts, we unexpectedly found that genes encoding components of the mating response MAPK signaling pathway were among the most significantly induced targets **(Fig. 3A, Table S2)**. This included strong upregulation of *STE2* (5.1 log_2_FC) and *STE4* (3.5 log_2_FC), which encode receptors for the α-factor mating pheromone, as well as *CEK2* (3.6 log_2_FC), encoding the terminal MAP kinase [31].

**Figure 3.**
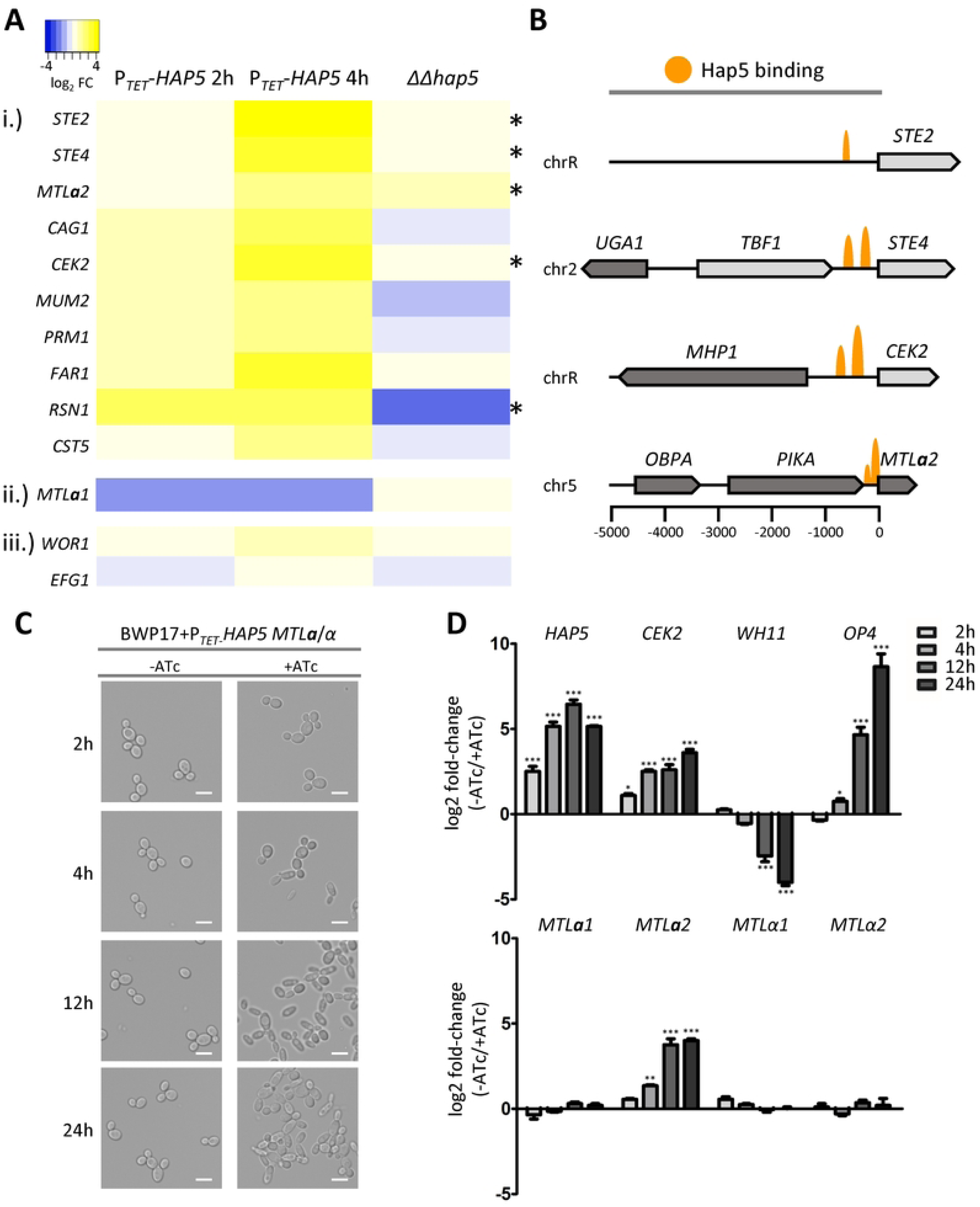
Regulation of mating-related genes by Hap5. **(A)** Heat map showing log2 fold-change levels of selected genes involved in mating (i), the gene encoding the repressor of mating (ii), and genes encoding master regulators of white-opaque switching (iii). Data represent transcript profiling of strains overexpressing *HAP5* (2 h and 4 h of induction) and *hap5* deletion mutants (ΔΔ*hap5*) grown in YPD medium at 30 °C. **(B)** Schematic overview of Hap5 binding peaks within selected promoter regions of identified target genes related to mating. **(C)** Cellular morphology of the control strain harboring the P*_TET_*-*HAP5* plasmid. Cells were grown in YPD liquid medium at **30 °C** for 24 h, either supplemented with anhydrotetracycline (+ATc; 3 µg/ml) or left untreated (-ATc). Representative microscopy images were taken at the indicated time points. Scale bars = 5 μm. **(D)** Transcript levels of *HAP5*, *CEK2*, the white-specific *WH11* gene, the opaque-specific *OP4* gene, and the mating-type master regulators (*MTL**a**1*, *MTL**a**2*, *MTLα1*, *MTLα2*) were determined by qRT-PCR. Transcript changes are expressed as log2 fold-change comparing induced (+ATc) to uninduced (-ATc) cells. Error bars represent ± SEM. Statistical significance was determined by a two-tailed Student’s *t*-test followed by a Bonferroni correction for multiple comparisons (* P<0.05; ** P<0.01; *** P<0.001).

In *C. albicans*, mating-related genes are strictly controlled by the **a**1-α2 repressor in *MTL**a**/α* heterozygous strains [6]. Intriguingly, despite our strains being *MTL**a**/α*, the *MTL**a**2* master regulator gene showed significantly increased expression upon *HAP5* overexpression. To rule out a loss-of-heterozygosity at the *MTL* locus, we performed genomic analytical PCR, which confirmed the presence of both *MTL**a**1* and *MTLα2* alleles in our strains **(Fig. S5)**. ChIP-seq data further revealed that Hap5 binds directly to the regulatory sequences of *STE2*, *STE4*, *CEK2*, and *MTL**a**2* **(Fig. 3B)**. These results indicate that Hap5 overexpression can overcome the canonical **a**1-α2-mediated repression to activate *MTL**a***-specific genes, yielding a distinct *MTL**a*** transcript signature in a heterozygous background.

### *HAP5*-overexpression enables white-to-opaque switching in *MTLa*/*α* heterozygous strains and their mating with *MTLa/a* diploids

Since the activation of mating pathways is intrinsically linked to the white-to-opaque phenotypic switch [2], we monitored cellular morphology following *HAP5* overexpression. While cells initially exhibited the classic white yeast shape, prolonged induction (12h to 24h) led to the appearance of elongated cells morphologically resembling opaque cells [32] **(Fig. 3C)**. This transition was confirmed via qRT-PCR, demonstrating a repression of the white-specific *WH11* gene and robust activation of the opaque-specific *OP4* gene [33,34] **(Fig. 3D)**.

To quantify this transition, we assessed white-to-opaque switching rates on solid media. Strains overexpressing *HAP5* formed typical pink opaque colonies on phloxine B-containing YP-K plates **(Fig. 4A)**. Interestingly, no opaque colonies were observed when these strains were grown on Lee’s medium supplemented with GlcNAc, a condition typically known to induce switching in *MTL*-homozygous strains, nor was switching ever observed in the empty-vector control under any condition **(Fig. 4A).** Because the CBC is tied to metabolism, we further tested the impact of nutrient availability. In agreement with recent findings that glucose starvation drives switching [8], glucose depletion drastically increased the switching rate of *HAP5*-overexpressing cells (up to ∼55%), whereas addition of glucose strongly repressed it **(Fig. 4B)**. Furthermore, because the CBC is a central regulator of iron homeostasis, we evaluated the effect of iron limitation. Strikingly, supplementing the medium with the iron chelator bathophenanthroline disulfonate (BPS) significantly reduced the Hap5-induced opaque switching rate (**Fig. 4C**).

**Figure 4.**
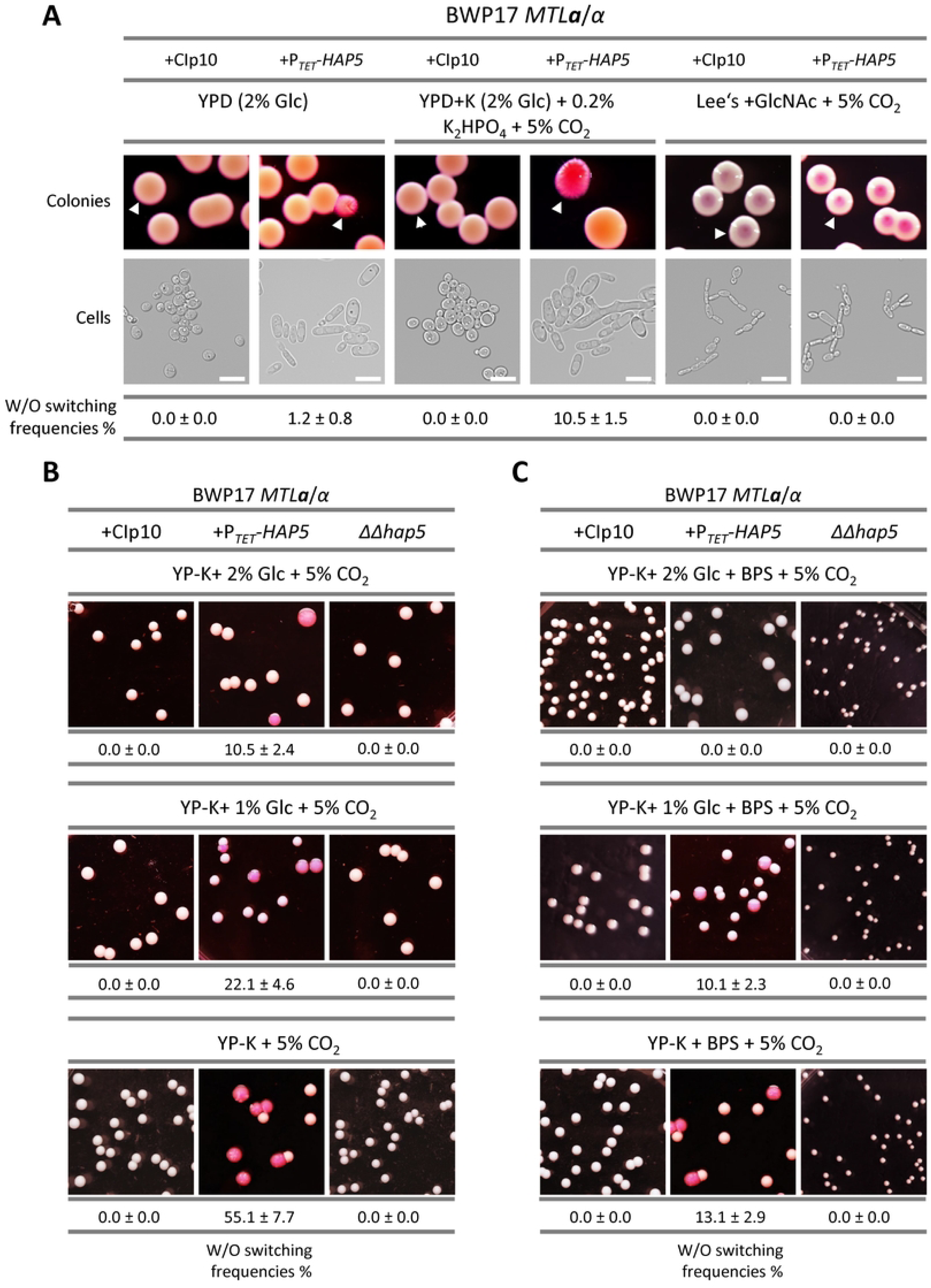
Cellular and colony morphologies of strains overexpressing *HAP5*. **(A)** Colony and cell morphology of *C. albicans* control strains harboring either the empty CIp10 plasmid or the P*_TET_*-*HAP5* plasmid. Strains were grown on YPD medium, YP-K (0.2% K_2_HPO_4_) + 5% CO₂, or Lee’s medium + 2% N-Acetylglucosamine + 5% CO₂, supplemented with phloxine B (5 µg/ml) and anhydrotetracycline (6 µg/ml) at 25 °C for 5 days. Cells were taken from the colonies indicated by a white triangle to be examined under the microscope. Scale bars = 5 μm. White-to-opaque switching frequencies were calculated as the percentage of red/pink colonies over total colonies (± SD). **(B)** Influence of glucose on switching. Strains were grown in a 5% CO₂ atmosphere on YP-K + 2% glucose, YP-K + 1% glucose, or carbohydrate-free YP-K medium (0% glucose) at 25°C for 5 days. **(C)** Influence of iron availability on switching. Strains were grown under the same conditions as in (B), but media were additionally supplemented with 5 mM of the iron-chelator BPS.

Crucially, to determine if these Hap5-induced opaque cells were mating-competent, we performed quantitative mating assays. *HAP5*-overexpressing *MTL**a**/α* cells successfully formed mating projections [6] and mated with a homozygous *MTL**a**/**a*** partner strain, whereas the control empty-vector strain did not **(Fig. 5)**. Remarkably, we also observed successful mating between two complementary *MTL***a**/α strains when both overexpressed *HAP5*, albeit at a lower frequency, demonstrating that metabolic rewiring via Hap5 is sufficient to unlock full sexual reproduction in ordinarily sterile heterozygous diploids.

**Figure 5.**
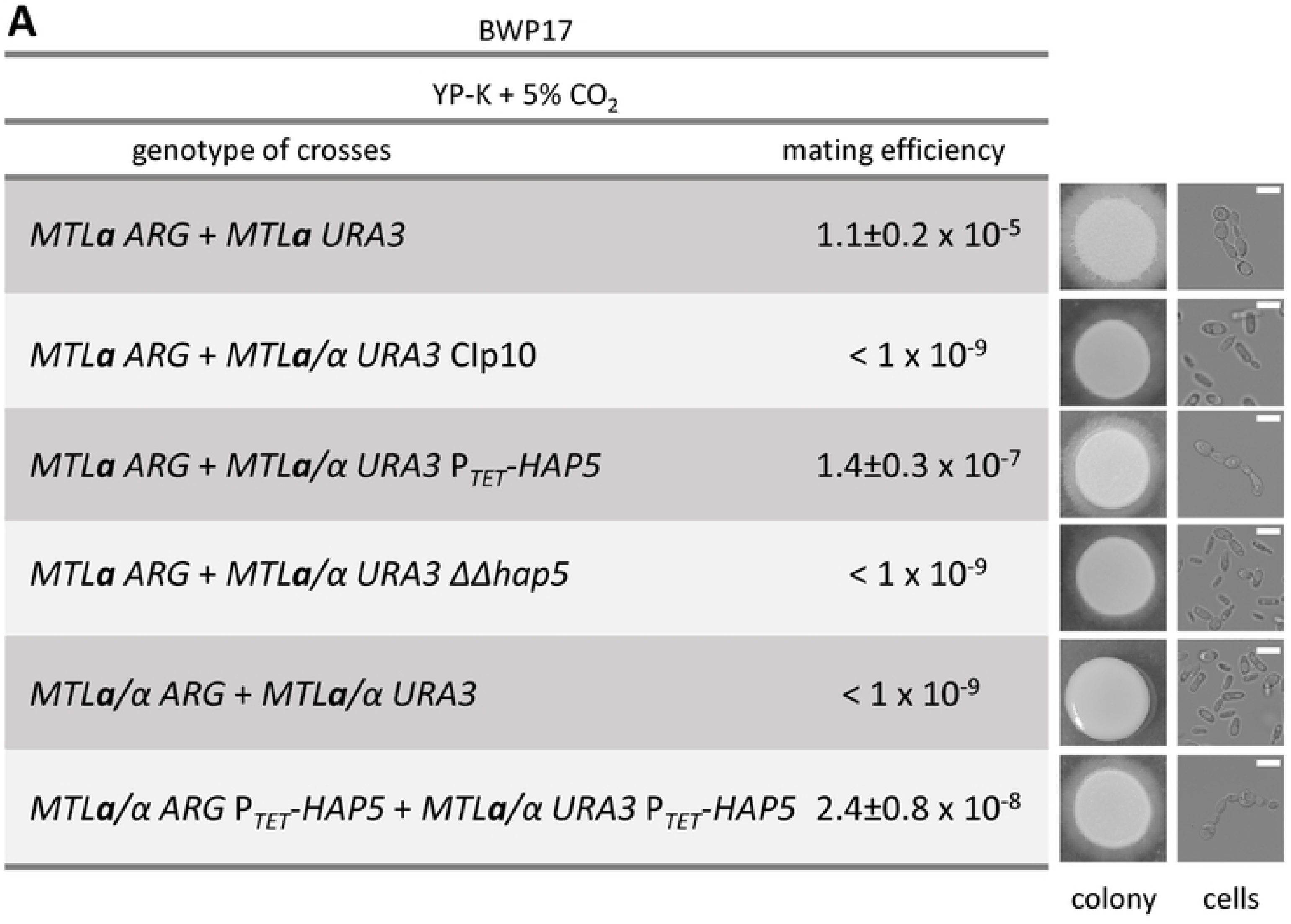
Overexpression of *HAP5* enables *MTL*a/α cells to undergo same-sex mating. Table representing the mating efficiency of the indicated crosses. Mixtures of indicated mating partners (2×10⁷ cells of each strain) were spotted and cultured on YP-K media in a 5% CO₂ atmosphere at 25 °C for 5 days, supplemented with anhydrotetracycline (+ATc; 6 µg/ml). Mating frequency was calculated as the number of colonies formed on selection plates relative to the total number of cells plated. A value of <1×10⁻⁹ indicates that no progeny was observed. Representative pictures of the resulting colonies and mating projections (cells) are shown on the right (scale bars = 3 μm).

### Identification of chromatin-associated proteins of the CCAAT-binding complex

To identify additional regulators that might play a role in this *HAP5*-driven white-to-opaque switching, we utilized our optimized ChIP-SICAP method to map the native protein interaction network of Hap5 at its genomic binding sites. Mass spectrometry identified 720 proteins enriched in the Hap5-TAP-tagged strain **(Fig. 6A, 6B; Fig. S1, Table S2)**. Network construction using PathoYeastract [25] revealed interactions with 19 transcription factors and 7 kinases **(Fig. 6C, 6D, 6E)**, mapping to diverse functions including filamentous growth (Flo8), carbohydrate metabolism (Tye7), and the oxidative stress response (Cap1) **(Table S4)**.

**Figure 6.**
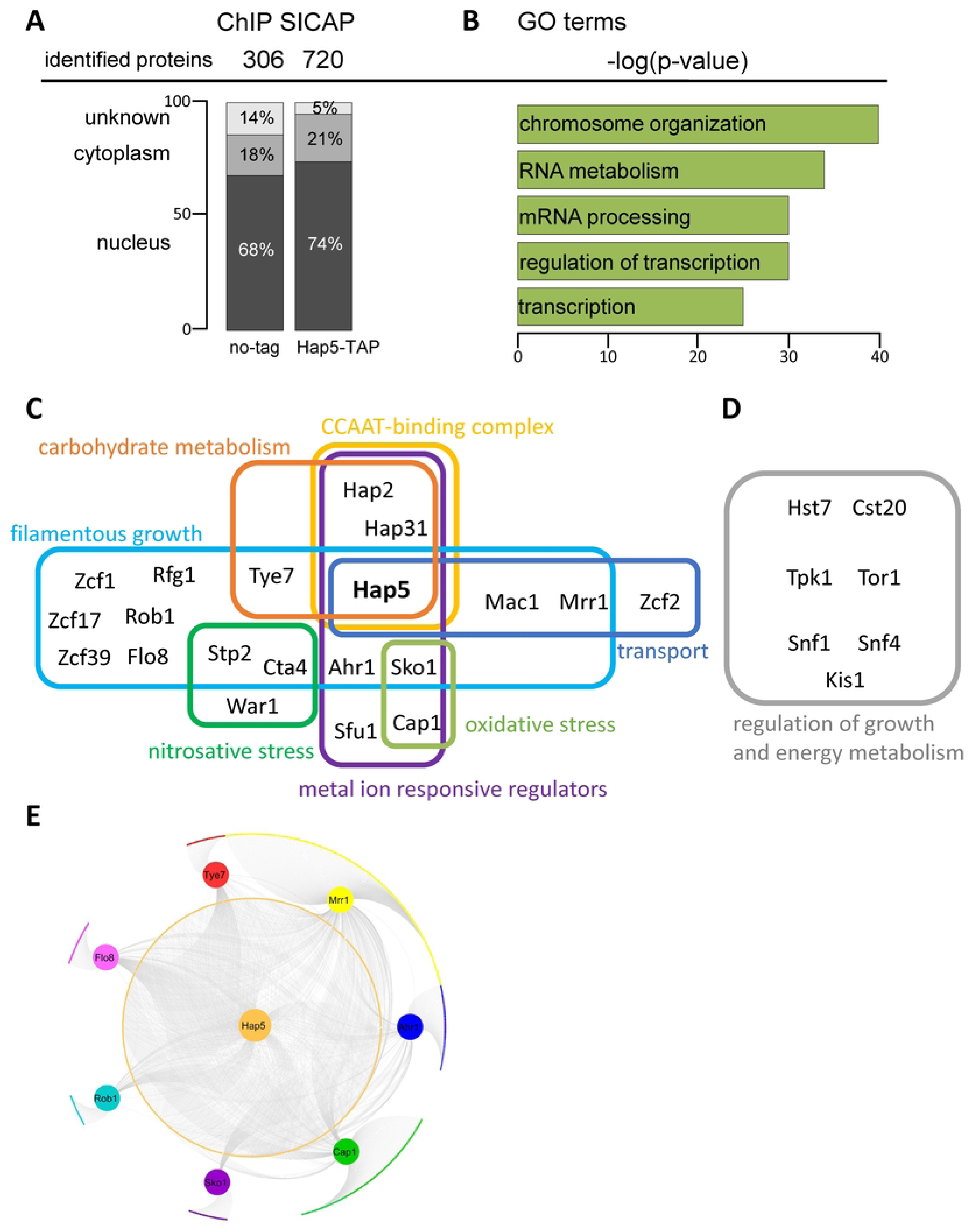
Interaction network of Hap5 identified by ChIP-SICAP. ChIP-SICAP (Selective Isolation of Chromatin-Associated Proteins) was used to identify potential interaction partners of TAP-modified Hap5. **(A)** Total number of identified proteins and their subcellular localization after immunoprecipitation in YPD medium at 30 °C. **(B)** Top 5 GO terms of identified proteins with corresponding p-values. **(C, D)** Identified transcriptional regulators (C) and kinases (D) categorized by their associated biological functions. **(E)** Visualized interaction network of Hap5 with potential regulators identified by ChIP-SICAP. Corresponding target genes of the respective transcriptional regulators were obtained from PathoYeastract and graphically displayed using Cytoscape.

### Cap1 is required for Hap5-driven white-to-opaque switching, whereas Hap43 acts antagonistically

To determine which of these newly identified interactors, as well as the known CBC components, were responsible for the mating phenotype, we performed epistasis analysis by integrating the *HAP5* overexpression construct into various transcription factor knockout backgrounds [35].

As expected, deletion of the essential core subunit *HAP2* [13] completely abolished *HAP5*-induced switching, while inactivation of *HAP31* or *HAP32* had no significant effect, confirming functional redundancy between these two subunits [14] **(Fig. 7)**. Interestingly, deletion of the iron-responsive co-regulator *HAP43* did not abolish the phenotype, but rather significantly *increased* the switching frequency (reaching ∼55%) **(Fig. 7)**. This indicates that Hap43 is not required for the induction of the mating program, but instead acts antagonistically in this context, likely competing for interaction with the Hap5 core complex to divert it toward iron acquisition rather than mating-type regulation.

**Figure 7.**
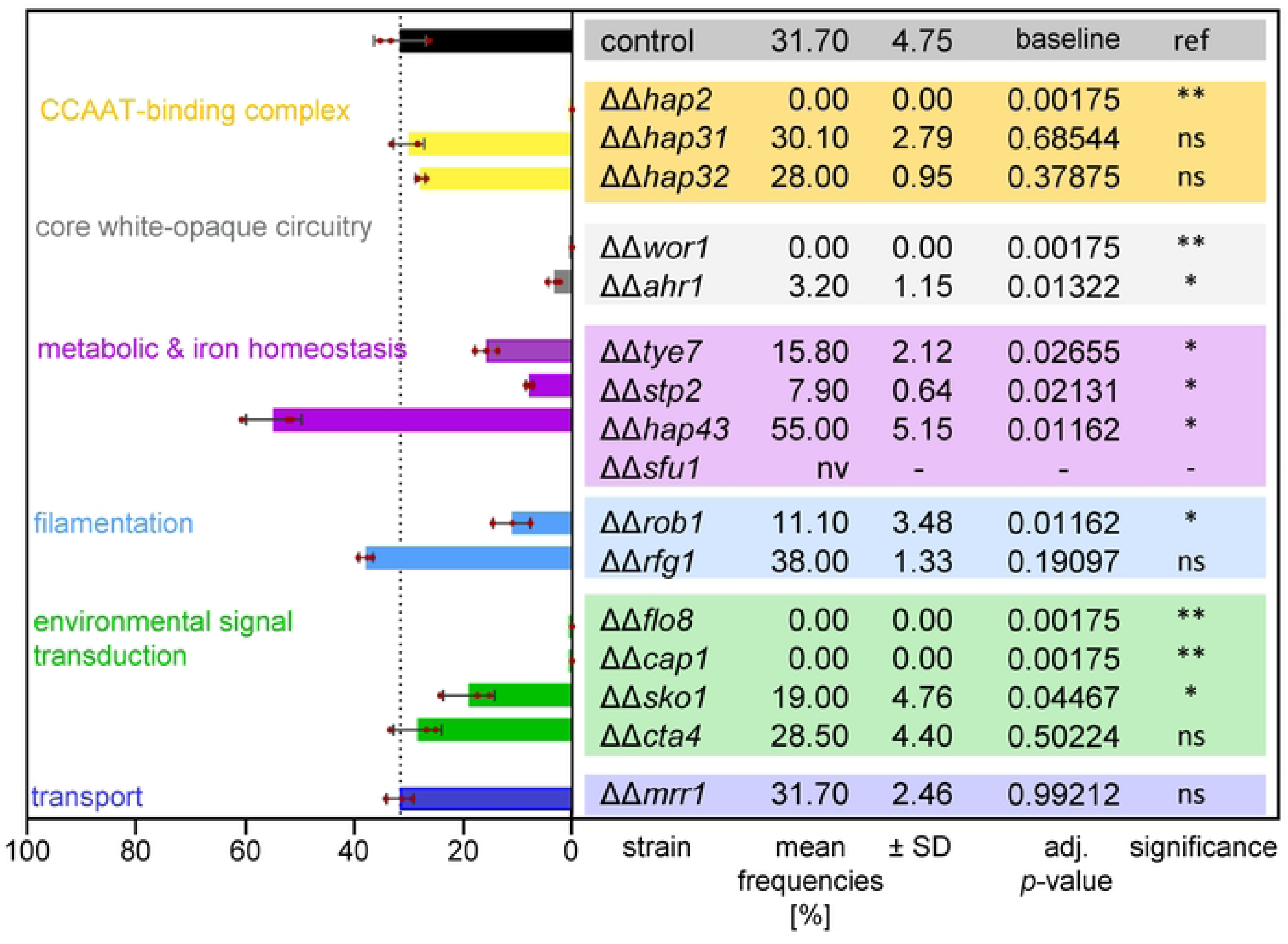
Quantitative epistasis analysis of the white-to-opaque switching network governed by Hap5. Absolute white-to-opaque switching frequencies of *C. albicans HAP5*-overexpression strains following the targeted homozygous deletion of specific downstream transcription factors. Mutants were sourced from the Homann transcription factor collection except for *wor1ΔΔ* [37], *flo8ΔΔ* [45] and *rfg1ΔΔ* (this study) and subsequently transformed with a P*_TET_*-*HAP5* overexpression plasmid. All were in a *MTL***a**/α background. Assays were performed on carbohydrate-free YP-K medium (0% glucose) with 6 µg/mL ATc following 5 days of incubation at 25 °C in a 5% CO₂ atmosphere. Horizontal bars represent the mean switching frequency derived from three independent biological replicates (n=3), with error bars indicating ± SD. The vertical dashed line marks the baseline switching capacity of the parental reference strain (control, 31.7%). Statistical significance was determined using a two-tailed Welch’s *t*-test comparing each deletion mutant directly against the parental reference, followed by a Benjamini-Hochberg FDR correction (* P<0.05; ** P<0.01; ns, non-significant; nv, non-viable).

Crucially, when we screened the newly identified Hap5-interactors, we found that deletion of general regulators like Flo8 and the mating master-regulator Wor1 [36,37] predictably abolished switching. Most importantly, the loss of Cap1 -the master regulator of the oxidative stress response -also fully abrogated the *HAP5*-induced white-to-opaque transition **(Fig. 7)**. This critical dependency suggested that the connection between Hap5 and mating competency is strictly routed through Cap1-mediated stress pathways.

### Cap1 interacts with Hap5

To physically validate this critical interaction, we performed co-immunoprecipitation between Hap5 and Cap1. Strains expressing TAP-tagged Hap5 and myc-tagged Cap1 were assayed under normal and iron-depleted conditions. Hap5 successfully co-precipitated with Cap1 under both growth conditions, unambiguously confirming their physical interaction at the chromatin level **(Fig. 8A)**. Detailed analysis of the Hap5-Cap1 co-regulatory network revealed a significant overlap in target genes governing energy production from carbohydrates and reactive oxygen species detoxification **(Fig. 8B)**, firmly linking metabolic status and oxidative stress to the developmental mating switch.

**Figure 8.**
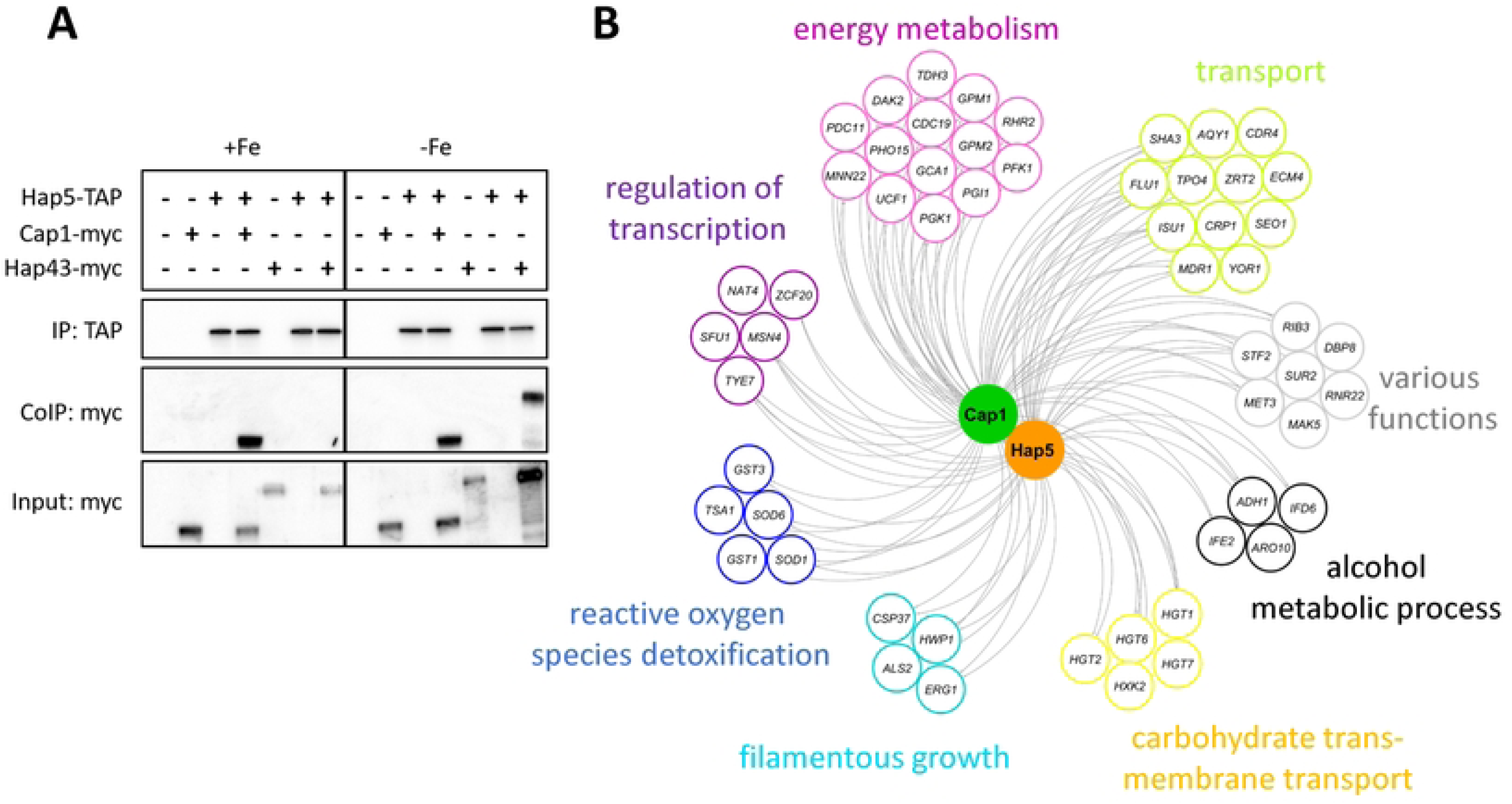
Cap1 interacts with Hap5. **(A)** Co-immunoprecipitation of Hap5 and Cap1. Strains overexpressing *HAP5*-TAP alongside either myc-modified Cap1 or Hap43 were grown in standard YPD medium (+Fe) or YPD medium treated with the iron chelator BPS (-Fe). The protein-A antigen of the TAP-tag was used for enrichment with IgG-coated magnetic beads. Immunoprecipitated samples were separated by SDS-PAGE, and western blot (WB) analysis was performed using an anti-myc antibody to detect Cap1-myc/Hap43-myc, and an anti-Protein A antibody to detect TAP-modified Hap5. Input samples were included as controls for myc-tagged protein levels. **(B)** The Hap5-Cap1 co-regulatory network. Common target genes of Hap5 and Cap1 were clustered based on their GO-term functions and visualized with Cytoscape.

## DISCUSSION

The ability of *Candida albicans* to undergo the white-to-opaque phenotypic transition is a fundamental developmental program that controls not only cellular morphology but also the capacity for sexual reproduction. Historically, mating in *C. albicans* has been viewed as a strictly controlled process requiring homozygosity at the mating-type locus (*MTL**a**/**a*** or *MTLα/α*), as the presence of the **a**1-α2 repressor in the vast majority of clinical heterozygous (*MTL**a**/α*) isolates tightly locks the cells in the sterile white state [6]. However, recent evidence suggests that extreme environmental pressures, such as glucose starvation, can override these genetic constraints and promote same-sex mating even in heterozygous populations [8]. Here, we identify the highly conserved CCAAT-binding complex (CBC) -with its core subunit Hap5 -as the missing mechanistic link that integrates nutritional and oxidative stress signals to bypass the **a**1-α2 repressor, effectively unlocking mating competency in typically sterile *MTL***a**/α diploids **(Fig. 9B)**.

**Figure 9.**
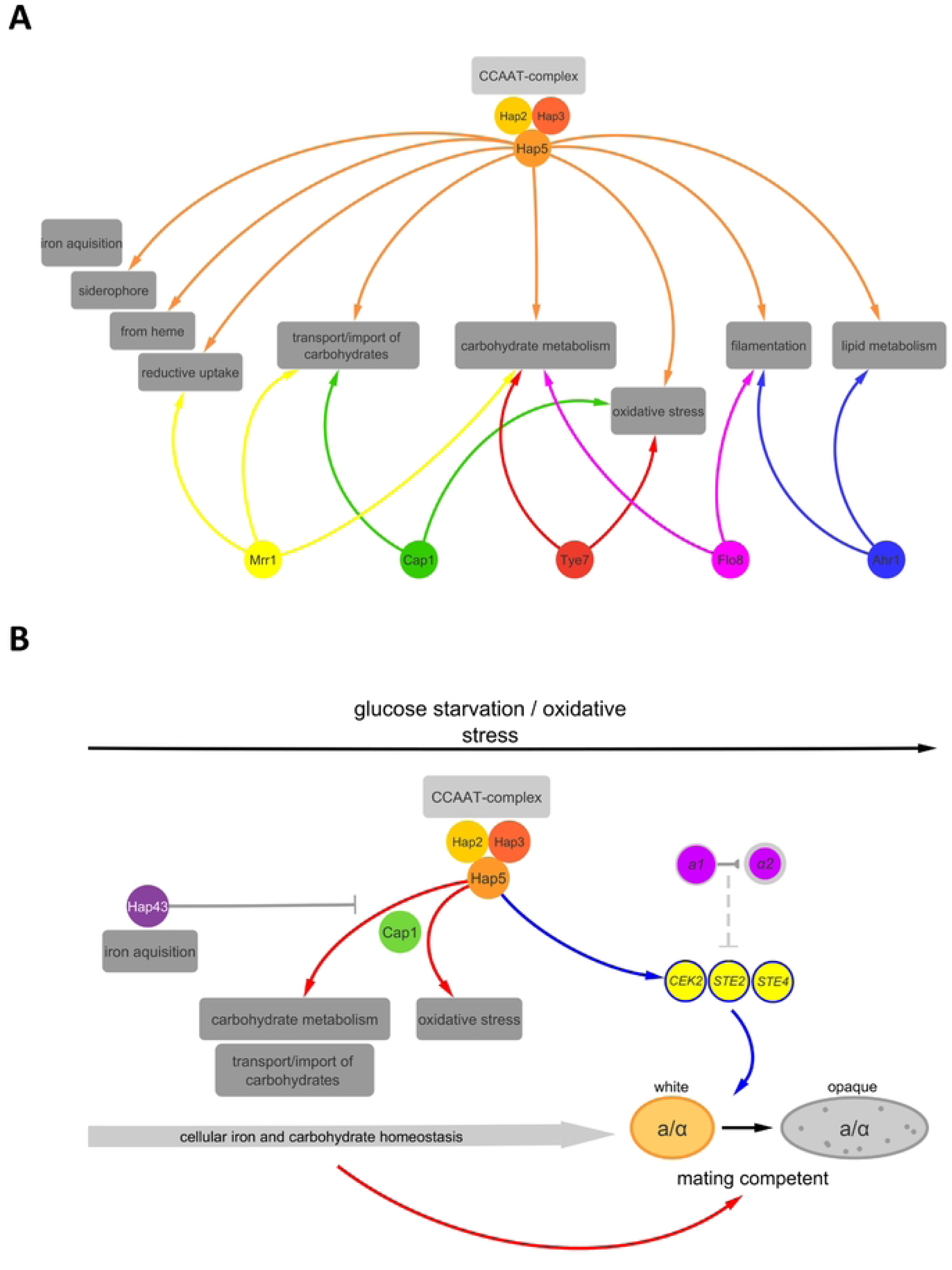
A unified model of CCAAT-complex functions and environmental adaptation. **(A)** Together with its identified co-regulators, Hap5 and the CCAAT-complex serve as a central regulatory hub governing iron acquisition, carbohydrate transport and metabolism, oxidative stress responses, filamentation, and lipid metabolism. **(B)** Model of Hap5-mediated white-to-opaque switch regulation. The CCAAT-transcription factor and Cap1 are jointly activated by harsh environmental signals (e.g., iron depletion, glucose starvation, oxidative stress). This complex directly binds to regulatory sequences of components of the mating signaling pathway (such as *CEK2*, *STE2*, and *STE4*). This activation overrides the standard **a**1-α2 repressor, enabling heterozygous *MTL***a**/α cells to become mating-competent and perform same-sex mating. This metabolic and stress-induced rewiring provides a competitive advantage and enables rapid adaptation to host environments.

Through combined transcriptomic and chromatin-binding analyses, we established that the CBC acts as a central hub for energy management, regulating genes involved in iron homeostasis, respiration, and carbohydrate metabolism [9] **(Fig. 9A)**. Strikingly, overexpression of *HAP5* induced a massive transcriptional rewiring that directly activated the mating MAPK signaling pathway (including *STE2*, *STE4*, and *CEK2*) and triggered the morphological transition to mating-competent opaque cells. We demonstrated that this pathway ultimately enables successful mating between genetically locked *MTL***a**/α strains.

### A unified model of environmental adaptation and mating competence

Taken together, our findings establish a comprehensive model for the regulatory scope of the CBC and how fungal pathogens overcome genetic sterility barriers (Fig. 9). Together with its identified co-regulators, Hap5 and the CBC serve as a central regulatory hub governing iron acquisition, carbohydrate transport and metabolism, oxidative stress responses, filamentation, and lipid metabolism (Fig. 9A). Crucially, we propose a model wherein the CBC and Cap1 are jointly activated by combinatorial hostile environmental signals, such as iron depletion, glucose starvation, and oxidative stress. This hyperactivated complex directly binds to regulatory sequences of components of the mating signaling pathway (such as *CEK2*, *STE2*, and *STE4*). This activation overrides the standard **a**1-α2 repressor, enabling heterozygous *MTL**a***/*α* cells to transition into the mating-competent opaque state and perform mating. Ultimately, this metabolic and stress-induced rewiring provides a competitive advantage and enables rapid adaptation to host environments (Fig. 9B).

The role of the CBC in governing major developmental transitions is not unprecedented. In the filamentous fungus *Aspergillus fumigatus*, the CBC acts as a central repressor of asexual reproduction (conidiation) in liquid cultures, a process that is reversibly regulated by environmental cues such as calcium stress [38]. Our finding that the *C. albicans* CBC controls the white-to-opaque transition extends this paradigm, demonstrating that the CBC serves as a deeply conserved developmental switch across diverse fungal lineages, repurposing its metabolic sensing capabilities to dictate cell fate and reproductive strategies.

A critical question arises: Does this switching occur under physiological conditions without artificial *HAP5* overexpression, and what environmental cues trigger it*?* A key caveat of our study is that all phenotypic readouts -including the mating transcriptomic signature, white-to-opaque switching, and mating competence -were observed under P*_TET_*-driven *HAP5* overexpression. Whether combinatorial host stresses activate endogenous Hap5 sufficiently to produce these outcomes remains to be tested directly. A second important limitation concerns the exact molecular mechanism by which Hap5 overcomes the *MTL* heterozygosity block. While bulk genotyping excluded genomic loss-of-heterozygosity, we did not directly assess **a**1 and α2 protein levels or their DNA occupancy during the transition. Therefore, it remains to be determined whether the activated CBC physically bypasses an actively bound **a**1-α2 repressor, or whether the extensive metabolic rewiring leads to the inactivation, destabilization, or displacement of the repressor complex. Notably, however, the Hap5-driven transition exhibits a highly specific metabolic dependency. While classic white-to-opaque switching is strongly induced by N-acetylglucosamine (GlcNAc) -a signal typically associated with the gastrointestinal tract or bacterial peptidoglycan [39] -*HAP5*-overexpressing cells failed to switch on GlcNAc-containing media. Instead, they strictly required glucose depletion. This differential response indicates that the hyperactivated CBC does not simply trigger a generic developmental program but acts as a highly specific sensor for distinct host niches. Our data suggest that the Hap5-driven mating program is tailored to the severe combinatorial stress of the host phagosome. When *C. albicans* is engulfed by host immune cells like macrophages or neutrophils, it encounters an environment characterized by exactly these conditions: severe nutrient scarcity (including glucose limitation) and intense oxidative bursts (Reactive Oxygen Species, ROS) [40,41].

Hap5 is constitutively expressed but functionally modulated by such environmental signals. Iron starvation necessitates the activity of the CBC for the acquisition of essential minerals [16], while ROS exposure serves as the direct, primary trigger for the activation and nuclear accumulation of the Cap1 transcription factor [42,43]. Our ChIP-SICAP and co-immunoprecipitation data revealed a direct physical and functional interaction between Hap5 and Cap1. The loss of *CAP1* completely abolished Hap5-mediated white-to-opaque switching, proving that their interaction is absolutely required to execute this developmental program. This metabolic interdependence highlights a sophisticated adaptive strategy: Hap5 regulates respiratory pathways that intrinsically generate endogenous ROS, while Cap1 handles the oxidative fallout. In the hostile environment of a phagosome, the simultaneous requirement for Hap5 (to adapt to iron/glucose starvation) and Cap1 (to survive oxidative bursts) creates a condition where both factors are highly active and interact at the chromatin level. We propose that it is this precise intersection of physiological stresses that physiologically triggers the Hap5-Cap1 mediated bypass of the *MTL* heterozygosity block **(Fig. 9B)**.

Furthermore, the dynamics of this regulatory hub are heavily influenced by competitive binding within the CBC. The iron-responsive factor Hap43 is known to interact with Hap5 to regulate iron acquisition [16]. Interestingly, we found that deletion of *HAP43* significantly exacerbated the white-to-opaque switching frequency. Conversely, inducing severe iron starvation with the chelator BPS actively suppressed the Hap5-driven switching rate **(Fig. 4C).** This mechanism of functional competition for the CBC core mirrors the regulatory architecture described in *C. glabrata*, where the accessory subunits Yap5 and Hap4 compete for CBC binding to dictate the transcriptional balance between iron tolerance and respiration, respectively [12]. In *C. albicans*, under exclusive iron starvation, Hap43 is strongly induced, dominates the complex, and drives iron acquisition while actively outcompeting the mating program. However, during combined oxidative and nutrient stress -or when Hap43 is absent -Cap1 can co-opt the CBC. This diversion effectively shifts the transcriptional machinery away from standard metabolic maintenance and toward a drastic developmental transition.

From an evolutionary perspective, unlocking sexual reproduction under severe combinatorial stress provides a distinct survival advantage. While mating is an energy-costly and time-consuming process, facilitating recombination in a typically sterile *MTL***a**/α population generates genetic diversity. In highly hostile environments where standard metabolic adaptations fall short, transitioning into an opaque state and engaging in mating may represent a “last resort” strategy to produce recombinant progeny better equipped to survive host defenses.

In conclusion, our findings identify the CBC as a candidate regulatory hub whose hyperactivation is sufficient to couple metabolic and stress sensing to a core developmental program, suggesting a potential mechanism for *C. albicans* to bypass genetic constraints on sexual reproduction that awaits validation under endogenous expression conditions.

## MATERIALS AND METHODS

### Strains, Plasmids, and Growth Conditions

*Candida albicans* strains used in this study are listed in **S1 Table**. Oligonucleotides used for strain construction are also provided in **S1 Table**. Unless otherwise stated, *C. albicans* cells were routinely cultured in YPD medium (1% (w/v) yeast extract, 2% (w/v) peptone, 2% (w/v) dextrose) at 30 °C. Where indicated, medium was supplemented with anhydrotetracycline (ATc; Sigma) at concentrations of 3 µg/mL (liquid cultures) or 6 µg/mL (solid media) to induce gene expression from the tetracycline-inducible P*_TET_* promoter. Solid media were prepared with 2% (w/v) agar.

Targeted gene knockout strains were obtained either from the Homann transcription factor collection [44] -including ΔΔ*hap2*, ΔΔ*hap31*, ΔΔ*hap32*, ΔΔ*hap43*, and ΔΔ*cap1*-or from other studies (*ΔΔflo8*, [45]; *ΔΔwor1*, [37]). Other mutants (e.g., ΔΔ*hap5* and ΔΔ*rfg1*) were generated *de novo* in the SN76 background using a transient CRISPR/Cas9 system [46]. Inducible *HAP5* overexpression strains were constructed by integrating either a pNIM1-derived [47] P*_TET_*-*HAP5* plasmid at the *ADH1* locus, or a CIp-P*_TET_-HAP5* plasmid [48] at the *RPS1* locus [19,20]. Strains expressing TAP-tagged Hap5 and untagged Hap5 controls were similarly generated. Constitutive overexpression was achieved in a SC5314 background deleted for *hap5* using the transient CRISPR/Cas9 system, transformed by a CIp-P*_TDH3_-HAP5* plasmid. Myc-tagged Cap1 and Hap43 strains were constructed in the wild-type background using the pFA-Clox system with 100 bp of homology flanking the respective genomic loci [49].

### ChIP-SICAP and LC-MS/MS Analysis

ChIP-SICAP (Selective Isolation of Chromatin-Associated Proteins) was performed as previously described [18], utilizing a protocol optimized for *C. albicans* [17]. Briefly, exponentially growing cells (OD₆₀₀ = 0.5) expressing either TAP-tagged Hap5 or untagged Hap5 (induced with 3 µg/mL ATc) were cross-linked with 1% formaldehyde for 15 minutes and quenched with 125 mM glycine. Chromatin was isolated and sheared to an average fragment size of 200–300 bp using a Bioruptor sonicator (Diagenode). Hap5-associated chromatin complexes were immunoprecipitated using IgG-coated magnetic beads. Following stringent washing, complexes were subjected to TEV protease cleavage (NEB). The retained chromatin was biotinylated using a chromatin-biotin labeling kit (Jena Bioscience) and further purified using streptavidin-coated beads (Thermo Fisher Scientific) followed by stringent washing to isolate exclusively chromatin-associated proteins. Proper enrichment and cleavage were validated by western blotting using an anti-TAP antibody (Fig. S1).

For mass spectrometry, chromatin-associated proteins on the streptavidin beads were subjected to on-bead trypsin digestion. Peptides were analyzed by LC-MS/MS using an Orbitrap Q-Exactive Plus mass spectrometer coupled to a Nano-LC Proxeon 1000 (Thermo Fisher Scientific). MS data were processed using MaxQuant, and proteins were identified by searching against the *C. albicans* proteome database. The analyses were performed in biological triplicates. To determine protein enrichment, LFQ intensities from the MaxQuant output were log2-transformed. Proteins were quantified if at least one unique peptide was identified. Reverse sequences, potential contaminants, and proteins identified only by site were discarded. For robust quantification, proteins were required to have at least two quantified intensity values per condition. Intensity values were then normalized by centering on the mean of the medians of intensities within each condition. Missing values in the dataset were imputed using the maximum likelihood estimation (impute.mle) function from the R package *imp4p*. Differential abundance analysis was performed using a LIMMA *t*-test. P-values were adjusted using an adaptive Benjamini-Hochberg correction, with the proportion of differentially abundant proteins estimated via the pounds method using the R package *cp4p*. Proteins were considered significantly enriched if they exhibited an absolute log_2_ fold-change > 2 and an adjusted p-value < 0.01. Additionally, proteins that were completely absent in the untagged control but consistently present in the TAP-tagged condition were directly classified as differentially abundant. The complete list of identified proteins, including LFQ intensities and statistical analyses, is provided in **S2 Table**. The mass spectrometry proteomics data have been deposited to the ProteomeXchange Consortium via the PRIDE partner repository with the dataset identifier PXD084794.

### Chromatin Recovery and ChIP-seq

To map Hap5 genomic binding sites, the chromatin retained on the trypsin-treated streptavidin beads was treated with an RNase mixture and incubated at 37 °C, followed by proteinase K and SDS treatment at 55 °C. Eluted DNA was purified and concentrated using AMPure XP beads. DNA integrity was evaluated on an Agilent Bioanalyzer High Sensitivity DNA chip. Libraries were constructed using the TruSeq ChIP Library Preparation Kit (Illumina) and sequenced on a NextSeq 550 system (Illumina). Reads were mapped to the indexed *C. albicans* genome using HISAT2. Peak calling was performed using the bPeaks program [21] and manually inspected using the Integrative Genomics Viewer (IGV; [50]). ChIP-MEME [23] was used for motif enrichment analysis. Network analyses were graphically displayed using Cytoscape [51], incorporating known target genes retrieved from PathoYeastract [52].

### RNA Sequencing (RNA-seq) and Data Analysis

To profile the Hap5 regulon, RNA-seq was performed on the *HAP5* overexpression strain (harvested 2 h and 4 h post-induction with ATc) and the *hap5* knockout mutant. Total RNA was extracted using the RNeasy Mini Kit (Qiagen). Libraries were prepared using the TruSeq Stranded mRNA Library Prep Kit (Illumina) and sequenced on a HiSeq 2500 system (Illumina). Transcriptomic data were processed using HISAT2, and differential gene expression analysis was conducted using DESeq2. RNA-seq data were submitted to the NCBI GEO under accession number GSEXXX (BioProject ID: XXXX).

### Reverse Transcription quantitative PCR (RT-qPCR)

Strains were cultured in YPD medium starting at an OD₆₀₀ of 0.1, either supplemented with 3 µg/mL ATc or left untreated. Cells were harvested at 2 h, 4 h, 12 h, and 24 h. Total RNA was isolated (RNeasy Mini Kit, Qiagen) and reverse-transcribed (QuantiTect, Qiagen). qRT-PCR was performed in MicroAmp Optical 96-well plates (Applied Biosystems) using the MX3000 Real-Time PCR system (Stratagene). Transcript levels were normalized to *ACT1* using the ΔΔCt method. Primers used for *HAP5*, *WH11*, *OP4*, *CEK2*, *MTL***a**1, *MTL*α1, *MTL***a**2, and *MTL*α2 are provided in **S1 Table**.

### Co-Immunoprecipitation (Co-IP) and Western Blotting

Strains co-expressing TAP-tagged Hap5 and myc-tagged Cap1 or Hap43 were grown in YPD medium with or without 5 mM of the iron chelator bathophenanthroline disulfonate (BPS) for 4 h. Cells were cross-linked with 1% formaldehyde for 15 min and quenched with 125 mM glycine. Cell pellets were lysed using glass beads in lysis buffer [50 mM HEPES-KOH pH 7.5, 140 mM NaCl, 1 mM EDTA, 1% Triton X-100, 0.1% sodium deoxycholate] supplemented with protease inhibitors. Cleared lysates were subjected to immunoprecipitation by incubation with IgG-coated magnetic beads (PanMouse IgG-coated, Thermo Fisher Scientific) for 4h at 4 °C. After extensive washing, eluted proteins were separated by SDS-PAGE and transferred to PVDF membranes. Total protein transfer was confirmed by membrane staining using a 0.1% Amido Black 10B solution (in 25% isopropanol and 10% acetic acid) for 5 min, followed by destaining. Membranes were blocked in 5% non-fat dry milk in TBST for 1h and protein interactions were detected by western blotting. TAP-tagged proteins were directly detected using an HRP-conjugated anti-TAP antibody (PAP, P1291, Sigma, diluted 1:1000). For the detection of myc-tagged proteins, membranes were incubated with a mouse monoclonal anti-myc primary antibody (clone 9E10, Abcam, diluted 1:1000), followed by incubation with an HRP-conjugated secondary antibody (diluted 1:10000). Signals were visualized using an enhanced chemiluminescence (ECL) system.

### White-to-Opaque Switching Assays

To assess white-to-opaque switching frequencies, overnight cultures of *MTL***a/**α heterozygous strains were diluted to an OD₆₀₀ of 0.1 in YP medium with varying glucose concentrations, supplemented or left untreated with 0.2% K₂HPO₄, or in Lee’s medium supplemented with 2% N-acetylglucosamine. Cultures were incubated for 2 days at 25 °C with gentle shaking. Cells were subsequently plated on the respective solid media containing 5 µg/mL phloxine B [32] and incubated for 5 days at 25 °C in a 5% CO_2_ atmosphere. Switching rates were calculated as the percentage of opaque (pink/red) colonies relative to the total number of colonies. Opaque cellular morphology was confirmed via microscopy.

### Mating Assays

Mating competency was assessed using *MTL***a**/α heterozygous strains harboring either the P*_TET_*-*HAP5* overexpression construct, the CIp10 empty vector, or the *hap5* deletion. Strains were incubated in YP-K medium supplemented with 3 µg/mL ATc at 25 °C for 24 h to induce opaque cell formation. Induced cells were collected and mixed with an opaque *MTL***a**/**a** mating partner strain in a 1:1 ratio (2 × 10⁷ cells each). The mixture was spotted onto glucose-depleted YP-K solid medium supplemented with 6 µg/mL ATc and incubated at 25 °C for 5 days. Mating frequency was calculated as the number of colonies formed on selection plates (SD medium lacking appropriate auxotrophic nutrients) relative to the total number of cells plated. Formation of mating projections was confirmed by microscopic inspection.

### Microscopy

Cells from liquid cultures or colonies on solid media were washed with PBS, transferred to microscope slides, and examined using a Leica DM RXA microscope equipped with a 100× oil-immersion objective (1.4 N/A). Macro images of colonies were taken from the bottom of the Petri dishes using a 40× objective or a digital camera and processed using ImageJ.

### Host Cell Damage and Enzymatic Assays

Host cell damage was evaluated using the oral keratinocyte cell line TR146 [53]. TR146 cells were grown to a confluent monolayer in DMEM supplemented with 10% FCS, penicillin, and streptomycin at 37 °C with 5% CO₂. 24 h prior to infection, the medium was replaced with Ham’s F12 medium (Gibco) containing 1% FCS. *C. albicans* yeast cells (2 × 10⁴) of the wild-type reference, the constitutive *HAP5* overexpression strain CEC5923, and the *hap5* knockout mutant CEC6078 were added to the TR146 cells and incubated for 24 h. Damage was quantified via lactate dehydrogenase (LDH) release using the Cytotoxicity Detection Kit (Roche/Sigma) according to the manufacturer’s instructions. Phospholipase and proteolytic activities were assessed by spotting 3 µL of a 10⁵ cells/mL suspension onto Egg Yolk agar and BSA agar, respectively, followed by incubation at 37 °C for 5 days. Enzymatic activity was determined by measuring the ratio of the colony diameter to the diameter of the precipitation zone.

### Use of AI-assisted technologies

During the preparation of this manuscript, the authors used Gemini Pro 3.1 (Google) to polish the English language and improve overall readability. After using this tool, the authors reviewed and edited the text comprehensively and take full responsibility for the final content of the publication.

### Data Availability

The mass spectrometry proteomics data (ChIP-SICAP) have been deposited to the ProteomeXchange Consortium via the PRIDE partner repository with the dataset identifier PXD084794 (Reviewer token: qKgHHXGPSmFL). The RNA-seq data generated in this study have been deposited in the NCBI Gene Expression Omnibus (GEO) under accession number GSE348730 (Reviewer token: iburmcgerrkbpmv). The ChIP-seq data generated in this study have been deposited in the NCBI GEO under accession number GSE348729 (Reviewer token: axcjaugqxvepfct).

## Acknowledgments

We thank Carlos R. Vázquez de Aldana for the generous gift of plasmids. We are grateful to Juliana Pipoli Da Fonseca for her support with the ChIP-seq and RNA-seq experiments at the Biomics Platform, and to Thibaut Douché at the Proteomics Core Facility for supporting the MS experiments (Institut Pasteur). We also thank Gaëlle Lelandais and Frédéric Devaux for sharing their expertise in NGS data analysis. Special thanks to Richard Bennett for numerous fruitful discussions during his stay in our laboratory, and for generously leaving piles of CHROMagar plates in our cold room. Finally, we thank all members of the Biologie et Pathogénicité Fongiques (BPF) unit for their continuous support and valuable insights.

## Author Contributions

**Conceptualization**

Lasse van Wijlick, Sophie Bachellier-Bassi, Christophe d’Enfert

**Methodology**

Lasse van Wijlick, Kontxi Martinez de San Vicente, Laxmi Shanker Rai, Natacha Sertour

**Investigation**

Lasse van Wijlick, Kontxi Martinez de San Vicente, Laxmi Shanker Rai, Natacha Sertour,

**Formal Analysis**

Lasse van Wijlick

**Writing – Original Draft Preparation**

Lasse van Wijlick

**Writing – Review & Editing**

Lasse van Wijlick, Sophie Bachellier-Bassi, Christophe d’Enfert

**Supervision**

Sophie Bachellier-Bassi, Salomé LeibundGut-Landmann, Christophe d’Enfert

**Funding Acquisition**

Salomé LeibundGut-Landmann, Christophe d’Enfert

## Supporting Information Legends

**S1 Table. Strains and oligonucleotides used in this study.**

**S2 Table. Comprehensive omics datasets for Hap5 (RNA-Seq, ChIP-Seq, and ChIP-SICAP).** This table contains three datasets: **(1) Transcriptome:** Differential gene expression analysis of the P*_TET_*-*HAP5* overexpression strain (after 2 h and 4 h of induction) and the *hap5* knockout mutant compared to the wild-type reference. **(2) ChIP-seq:** Genome-wide binding coordinates and annotated target genes of Hap5. **(3) ChIP-SICAP (LC-MS/MS):** Proteomic analysis of the Hap5 chromatin-associated interaction network. The dataset includes Uniprot accessions, gene names, the number of identified peptides, and molecular weights [kDa]. Abundance is provided as individual LFQ intensity scores across biological triplicates for both the untagged control (*HAP5_1*, *HAP5_2*, *HAP5_3*) and the tagged strain (*HAP5-TAP_1*, *HAP5-TAP_2*, *HAP5-TAP_3*). Statistical enrichment is detailed via log2FoldChange, p-value, adjusted p-value (p-adj), and a significance cut-off indicator (FDR < 1%).

**S3 Table. GO-term analysis results of the Hap5-network.**

**S4 Table. GO-term analysis results of the Hap5 co-regulatory network.**

**Figure S1. Validation of the Hap5-TAP tagging construct and ChIP-SICAP enrichment. (A)** Schematic representation of the vector used to overexpress TAP-tagged Hap5 from a tetracycline-inducible promoter. The construct allows for C-terminal TAP tagging, with the TAP tag (approx. 20 kDa) being composed of a Calmodulin Binding Peptide (CBP) and a Protein A (ProtA, approx. 15 kDa) antigen separated by a TEV cleavage site. **(B)** Formaldehyde-crosslinked Hap5-TAP from three biological replicates used for ChIP-SICAP was separated by SDS-PAGE and analyzed by western blot with an anti-TAP antibody. TEV cleavage activity was monitored over time, demonstrating that 4 h is sufficient to release Hap5-CBP (indicated by membrane staining).

**Figure S2. Comparative transcriptomics of iron-responsive genes regulated by Hap5.** Venn diagrams showing the overlap of transcript profiles between cells grown under iron-depleted conditions [27] and strains overexpressing *HAP5* after **(i)** 2 h and **(ii)** 4 h of induction, as well as **(iii)** the *hap5* deletion mutant (ΔΔ*hap5*).

**Figure S3. Extended comparison of HAP5 transcript profiles.** Venn diagrams detailing the overlap of differentially expressed genes across different conditions. **(A)** Overlap of *HAP5* overexpression profiles after 2 h and 4 h of induction. **(B, C)** Comparison of the *HAP5* overexpression profile after 2 h (B) and 4 h (C) with the transcript profile of the *hap5* deletion mutant. **(D)** Identification of commonly upregulated and downregulated genes across both time points of *HAP5* overexpression compared to the *hap5* deletion mutant.

**Figure S4. Regulation of virulence effectors by Hap5. (A)** Heat map showing log2 fold-change levels of genes encoding secreted aspartyl proteinases (*SAPs*) and the candidalysin toxin (*ECE1*). Data represent transcript profiles of strains overexpressing *HAP5* (2 h and 4 h) and the *hap5* deletion mutant grown in YPD medium at 30 °C. **(B-D)** Phenotypic virulence assays using a strain constitutively overexpressing *HAP5* (P*_TDH3_*-*HAP5*) and the *hap5* knockout mutant. Assays show **(B)** damage induced in human oral epithelial (TR146) cells quantified by lactate dehydrogenase (LDH) release, **(C)** proteolytic activity on BSA-containing agar plates, and **(D)** phospholipase activity on Egg Yolk-containing agar plates. Statistical significance is indicated (* P<0.05; ** P<0.01; **** P<0.0001).

**Figure S5. Verification of the *MTL* genotype in white and opaque cells overexpressing *HAP5*.** Analytical PCR was used to determine the *MTL* genotype of cells that either remained white (W) or switched to opaque (OP) after isolation on YP-K medium containing anhydrotetracycline (6 µg/mL). Genomic DNA was analyzed by PCR using oligonucleotides specific for either *MTL***a** or *MTL*α and separated on a 1% agarose gel. The heterozygous parental strain (CEC2907), a hemizygous *MTL**a**/Δ* strain (CEC2257), and a hemizygous *MTLα/Δ* strain (CEC2264) were included as controls, confirming that all tested *HAP5*-overexpression strains maintained their *MTL**a**/α* heterozygosity.

